# Mutations in *Drosophila* tRNA processing factors cause phenotypes similar to Pontocerebellar Hypoplasia

**DOI:** 10.1101/2021.07.09.451847

**Authors:** Casey A. Schmidt, Lucy Y. Min, Michelle H. McVay, Joseph D. Giusto, John C. Brown, Harmony R. Salzler, A. Gregory Matera

## Abstract

Mature tRNAs are generated by multiple RNA processing events, which can include the excision of intervening sequences. The tRNA splicing endonuclease (TSEN) complex is responsible for cleaving these intron-containing pre-tRNA transcripts. In humans, TSEN copurifies with CLP1, an RNA kinase. Despite extensive work on CLP1, its *in vivo* connection to tRNA splicing remains unclear. Interestingly, mutations in *CLP1* or *TSEN* genes cause neurological diseases in humans that are collectively termed Pontocerebellar Hypoplasia (PCH). In mice, loss of Clp1 kinase activity results in premature death, microcephaly and progressive loss of motor function. To determine if similar phenotypes are observed in *Drosophila*, we characterized mutations in *crowded-by-cid* (*cbc*), the *CLP1* ortholog, as well as in the fly ortholog of human *TSEN54*. Analyses of organismal viability, larval locomotion and brain size revealed that mutations in both *cbc* and *Tsen54* phenocopy those in mammals in several details. In addition to an overall reduction in brain lobe size, we also found increased cell death in mutant larval brains. Ubiquitous or tissue-specific knockdown of *cbc* in neurons and muscles reduced viability and locomotor function. These findings indicate that we can successfully model PCH in a genetically-tractable invertebrate.

## Introduction

Transfer (t)RNAs play a crucial role in the heavily regulated process of protein expression. As highly structured adaptors, tRNAs translate nucleic acid messages into polypeptide outputs. In many organisms, including metazoans, most tRNA genes need only to be transcribed, end-processed, and modified before they can participate in protein translation (1). However, a subset of tRNA genes contain introns (2,3). These intervening sequences are generally small and interrupt the anticodon loop of the tRNA (4,5). Because tRNA structure is crucial for proper function, intron removal is a critically important step in tRNA biogenesis (5).

Metazoan tRNA intron removal is carried out in two enzymatic steps. First, a heterotetrameric enzyme complex, called TSEN, recognizes the pre-tRNA transcript (6). This complex cleaves the pre-tRNA in two places: at the 5′ exon-intron boundary, and at the intron-3′ exon boundary (7). These cleavage events produce 2′,3′-cyclic phosphates on the 3′ ends of the 5′ exon and intron, and 5′-OH on the 5′ ends of the intron and 3′ exon (8). Once the intron has been cut out of the pre-tRNA, a ligase enzyme, called RtcB, joins the two exon halves together using the cyclic phosphate on the 5′ exon as the junction phosphate to generate a mature tRNA (9). Strikingly, RtcB can only use the 5′-OH on the 3′ exon as a substrate for ligation; it cannot use a 5′-phosphate (9). In human cells, RtcB functions in a complex with other factors including archease and DDX1 (10); archease co-puriefies with RtcB and cooperates with DDX1 to stimulate full activity of RtcB in biochemical assays (10). For many years, the fate of the excised tRNA intron was unknown. However, recent work from our lab has shown that *Drosophila* tRNA intron ends are ligated together to yield a unique subspecies of circular RNA, called a tRNA intronic circular RNA or tricRNA (5,11,12). Earlier reports show that circularized tRNA introns (tricRNAs) from several archaeal species are generated in a similar manner (13). Because this ligation event also utilizes the 2′,3′-cyclic phosphate as the junction phosphate, the overall process is termed the “direct ligation” tRNA splicing pathway (12).

In contrast to direct ligation, plants and fungi utilize the “healing and sealing” ligation pathway (4). Pre-tRNA cleavage is carried out by orthologs of the TSEN complex and the same non-canonical RNA ends are generated. However, the method of ligation is quite distinct. A multifunctional enzyme called Rlg1/Trl1 executes three activities: the 5′-OH of both the intron and 3′ exon are phosphorylated via its kinase domain; the 2′,3′-cyclic phosphate of the 5′ exon is opened through cyclic phosphodiesterase activity; and the exon halves are then joined by the ligase domain. The extra phosphate at the junction of the newly ligated tRNA is removed by a 2′-phosphotransferase enzyme called Tpt1 (14). In contrast to the pathway employed by archaea and metazoa, the tRNA intron, now containing a 5′-phosphate, becomes a substrate for degradation by the exonuclease Xrn1 (15).

Although there has been an extensive body of work on tRNA splicing in yeast and human cell culture models, less is known about the function of metazoan tRNA splicing enzymes *in vivo*. Among the many human neurological diseases, there is a class of disorders called Pontocerebellar Hypoplasia (PCH). Interestingly, several subytpes of PCH are associated with mutations in genes that encode the TSEN complex (16–22). Although the precise disease mechanisms are not well understood, each PCH subtype exhibits structural abnormalities in the brain, including microcephaly. These growth and tissue maintenance defects lead to developmental delays, mobility issues, and intellectual disabilities. Consistent with these reports in humans, RNAi-mediated knockdown of *Tsen54* in zebrafish embryos was shown to cause structural defects as well as cell death within the brain (23). Further experimentation showed that genetic knockout of *Tsen54* was lethal in these animals (23).

Recently, a new subtype of PCH was identified in several consanguineous families from eastern Turkey (24–26). PCH10 is associated with missense mutations in the human *CLP1* polyribonucleotide 5′-hydroxyl kinase gene (27). Originally identified as part of the mRNA 3′ end processing machinery (28), CLP1 was subsequently found to copurify with the human TSEN complex, suggesting a role for this protein in pre-tRNA processing (6). Indeed, CLP1 can phosphorylate the 5′-ends of tRNA 3′ exons *in vitro* (27). This finding was remarkable, considering that phosphorylation of the 5′-OH is known to inhibit RtcB-mediated ligation (9). We previously demonstrated that CLP1 is neither required for tRNA intron cleavage, nor for tRNA exon ligation (29). Instead, CLP1 kinase activity serves as a negative regulator of tRNA processing, perturbing exon ligation as well as intron circularization (29). Strikingly, all of the PCH10 cases reported to date document the same homozygous *CLP1* missense mutation: a G>A transition resulting in substitution of a histidine for an arginine residue (R140H) (24–26). This mutation has been shown to reduce the RNA kinase activity of CLP1 as well as its ability to associate with the TSEN complex (24,25). Importantly, a Clp1 kinase-dead mouse model displayed similar neurodegenerative phenotypes to those of human PCH10 patients (30).

Although tRNA disease-related phenotypes have been characterized in several vertebrate models, much less is known about tRNA processing in these systems, particularly the *in vivo* fate and status of tRNA introns. Due to the small size of vertebrate tRNA introns, most of which are less than 25nt long (3), they are difficult to detect. Given that human tRNA introns are very likely circularized (31), they would therefore evade typical sRNA sequencing approaches that depend on adaptor ligation prior to reverse transcription. These features make it difficult to study certain aspects of tRNA processing defects in vertebrate systems *in vivo*.

In contrast, the introns of several invertebrate tRNA genes are comparatively large (11). For example, one *Drosophila* tRNA gene (*CR31905*) contains a 113nt intron, and its resulting tricRNA can be easily detected throughout development by northern blotting and RNA sequencing (11). In addition, its topology (i.e., linear vs. circular) can be verified by molecular biological assays (11). We also previously characterized tRNA processing factors in *Drosophila* cells, including Tsen54 (12), and the fly ortholog of CLP1, crowded-by-cid (cbc) (29). Furthermore, there are sophisticated genetic tools available in flies that allow manipulation of gene expression in a very fine tissue-specific or developmental time-specific manner. In this study, we establish a *Drosophila* model of PCH by characterizing available *Tsen54* and *cbc* mutants. We show that these animals exhibit viability, locomotor, and brain size phenotypes similar to those observed in PCH patients and vertebrate models. We also use RNA interference (RNAi) to determine if there is a tissue-specific requirement for cbc expression. In summary, this work establishes a new animal model of PCH wherein one can more readily examine tRNA splicing outputs.

## Methods

### Fly lines and husbandry

We obtained *Tsen54* and *cbc* mutant alleles from the Bloomington stock center (Table 1). The balancers for the *cbc* and *Tsen54* mutant lines were changed as follows: for the *cbc* alleles, the balancer was switched with the CyO-Actin-GFP second chromosome balancer; for the *Tsen54* alleles, the balancer was switched with the TM6B-GFP third chromosome balancer.

**Table 1.** *Drosophila* mutants used in this study.

| Human gene | Fly gene | Bloomington stock number | Mutation |
| --- | --- | --- | --- |
| CLP1 | cbc | 27620 | point mutant; A37T |
| CLP1 | cbc | 27621 | point mutant; Q5X |
| CLP1 | cbc | 17812 | p-element insertion in the 2nd exon |
| TSEN54 | Tsen54 | 18077 | p-element insertion in the 3' UTR |
| TSEN54 | Tsen54 | 36235 | p-element insertion in the 4th exon |

We obtained RNAi lines from either the Bloomington or Vienna stock centers (Table 2). Stock numbers that begin with ‘v’ are from Vienna.

**Table 2.** RNAi lines used in this study.

| **RNAi line** | **Stock number** |
| --- | --- |
| TSEN2 RNAi | 55659 |
| TSEN54 RNAi #1 | 35337 |
| TSEN54 RNAi #2 | 57756 |
| RtcB RNAi #1 | v36198 |
| RtcB RNAi #2 | v36494 |
| Ddx1 RNAi #1 | 55744 |
| Ddx1 RNAi #2 | 27531 |
| Archease RNAi | 42851 |
| CG33057 RNAi #1 | 55699 |
| CG33057 RNAi #2 | 62962 |

The Gal4 drivers used in this study (Table 3) were obtained from Dr. Ashlyn Spring.

**Table 3.** Gal4 drivers used in this study.

| **Gal4 driver** | **Affected tissue** |
| --- | --- |
| daughterless | Ubiquitous |
| GMR | Eye |
| C15 | Neurons + muscle |
| C155 | Neurons |
| C164 | Motor neurons |
| OK371 | Glutamatergic neurons |
| Cha | Cholinergic neurons |
| C57 | Muscle |
| Mef2 | Muscle |

Stocks were maintained on molasses food. All fly stocks and crosses were performed at 25°, except for RNAi crosses which were performed at 29°. To generate homozygous mutant animals, males and females from a balanced stock were placed in a cage to mate, and females laid eggs on molasses plates with yeast paste. After allowing the plates to age for one day, GFP-negative larvae were sorted into vials for viability and locomotion assays (so as to reduce competition from heterozygous siblings). Rescue genotypes were generated in the same manner as mutant animals, except females laid eggs on grape juice plates rather than molasses plates.

### Viability assay

The viability assay was performed as described in (32).

### Locomotor assay

The larval locomotion assay was performed as described in (32) with the following modifications: videos were taken on an iPhone 8 and trimmed to exactly 45 seconds.

### Brain dissection and imaging

Third instar larvae were gross dissected in PBS using sharpened forceps to expose the larval brain. The inverted carcasses were fixed in 4% formaldehyde, washed three times in PBT (0.1% Triton X-100 in PBS), blocked in PBT-G (1% normal goat serum + 0.3% Triton X-100 in PBS) and incubated in primary antibody diluted in PBT-G overnight at 4°. The next day, the inverted carcasses were washed three times in PBT, incubated in secondary antibody + Phalloidin-FITC for one hour at 25°, and washed three additional times in PBS before fine dissecting the larval brains. The brains were mounted on a slide with mounting media containing DAPI. The slides were imaged using a Zeiss 710 confocal microscope, and the images were processed with ImageJ. The following antibodies were used in this study: rabbit anti-cleaved caspase 3 D175 at 1:100 (Cell Signaling Technology #9661); Alexa 647 goat anti-HRP at 1:500 (Jackson ImmunoResearch Laboratories Inc. (123-605-021); Alexa 488 goat anti-rabbit at 1:1000 (Life Technologies ref# A11008).

## Results

Mutant lines used in this study were obtained from the Bloomington Drosophila Stock Center. A list of the alleles we obtained is shown in Table 1. As shown in Figure 1A, the *cbc*^1^ and *cbc*^2^ alleles are EMS-induced mutations generated by the Kaufman lab that had been deposited at Bloomington, but never characterized (33). The *cbc*^3^ allele is a p-element insertion in the *cbc* 3′ UTR. We also obtained two p-element insertion alleles for *Tsen54*, called *Tsen*54^1^ and *Tsen54*^2^, corresponding to insertions in the second and fourth exon, respectively (Fig. 1B). Using these stocks, we set out to establish a tRNA splicing-based model of PCH in *Drosophila*.

**Fig 1.**
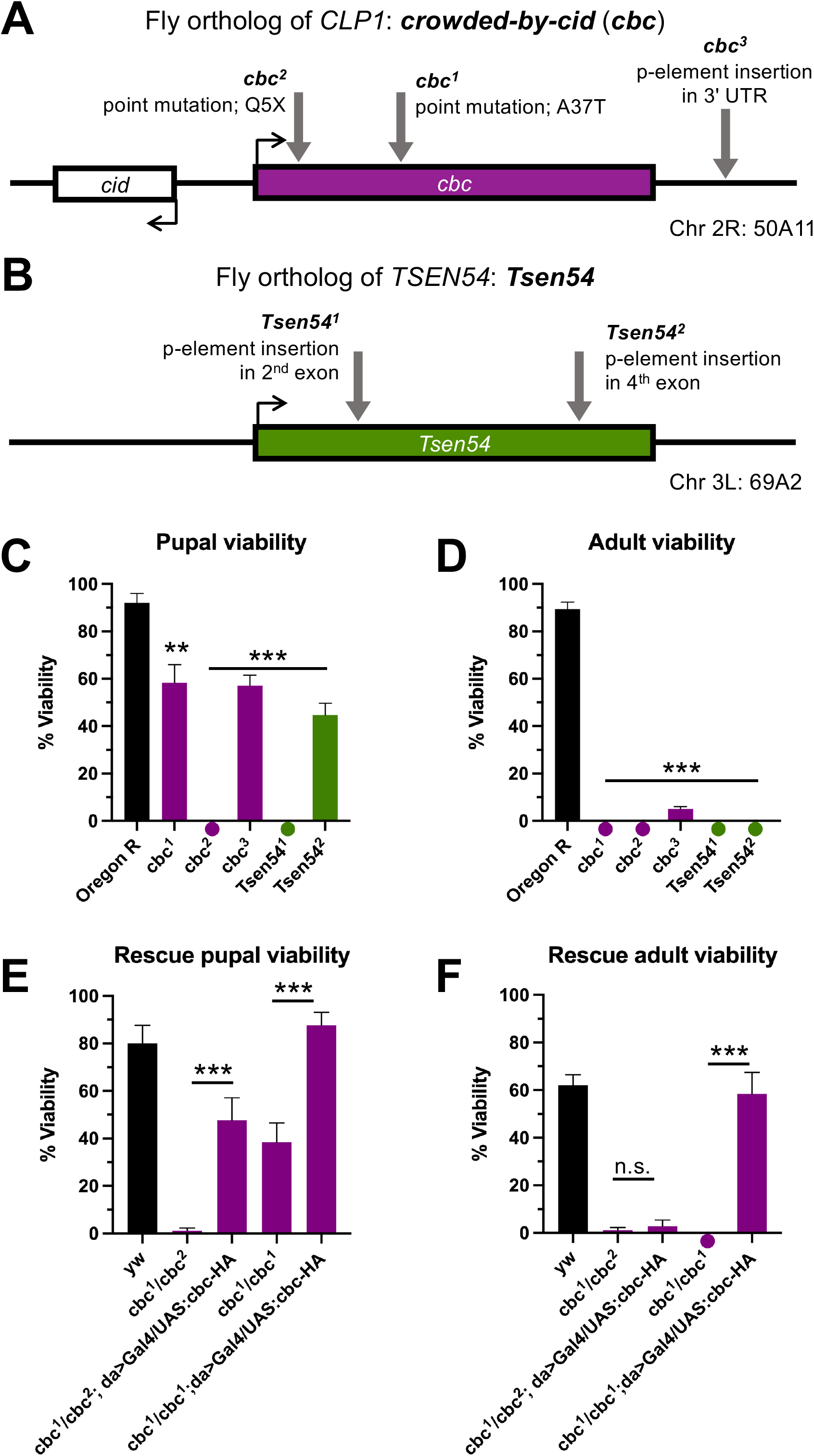
Mutations in *Drosophila* tRNA processing factors cause strong viability defects. (A) and (B) Summary of mutant alleles obtained from the Bloomington Drosophila Stock Center for *cbc* (A) and *Tsen54* (B). (C) Pupal and (D) adult viability of homozygous mutants (*cbc* in purple, *Tsen54* in green) compared to control (Oregon R in black). (E) Pupal and (F) adult viability of rescue crosses compared to control (yw in black). Note that in some mutants a circle is placed below the x-axis to indicate a zero value rather than absence of data. ** p<0.01; *** p<0.001, student’s t-test. n.s., not significant.

### tRNA processing factor mutants display severe viability defects

We first examined viability in homozygous tRNA processing factor mutants using a standard viability assay. As shown in Fig. 1C, we identified significant defects in each of the mutants tested. Notably, the *cbc*^2^ allele was quite severe; these animals all die as embryos. Similarly, the *Tsen54*^1^ homozygotes die as third instar larvae. The remaining mutants (*cbc*^1^, *cbc*^3^, and *Tsen54*^2^) display a pupation frequency of ~50% (Fig. 1C), and only the *cbc*^3^ homozygotes eclose as adults (albeit at a very low frequency, ~5%; Fig. 1D).

We next tested whether the viability defects observed in the cbc homozygous mutants could be rescued by expression of wild-type *cbc*. We ubiquitously expressed full-length, HA-tagged cbc (34) in the background of both *cbc*^1^ homozygous mutants and *cbc*^1^/*cbc*^2^ trans-heterozygotes. Expression of wild-type *cbc* significantly rescued pupal viability in both the homozygous and trans-heterozygous mutant backgrounds (Fig. 1E), and adult viability in the homozygous mutants (Fig. 1F).

These results are consistent with human PCH patient data, where one study reported the median age of death to be 50 months (17). Additionally, the observed viability defects are recapitulated in animal models of PCH. For example, zebrafish bearing a homozygous nonsense mutation in *Tsen54* did not survive beyond 10 days post-fertilization (23). Furthermore, *Clp1*-null mice die before embryonic day 6.5, a very early time point in mouse development (30). We conclude that homozygous mutations in tRNA processing factors cause strong viability defects in *Drosophila*.

We also assessed viability in animals depleted of known *Drosophila* tRNA processing factors (12) using RNA interference (RNAi) via the Gal4-UAS system (35). We obtained UAS-RNAi lines for *Drosophila* tRNA processing factors (Table 2) and crossed these animals to either Oregon R (to control for the UAS-RNAi transgene) or to daughterless-Gal4 (to ubiquitously express shRNAs against the targeted tRNA processing factor). Additional controls for this experiment were Oregon R (wild-type) and daughterless-Gal4 alone. None of the control crosses displayed pupal (data not shown) or adult (Fig. 2, black bars) viability defects. Interestingly, ubiquitous depletion of TSEN2, TSEN54, RtcB, and Ddx1 had no effect on pupation efficiency; however, animals expressing shRNA against archease failed to pupate and died as larvae (data not shown). RNAi knockdown of TSEN2 or TSEN54 did not greatly affect adult viability (Fig. 2, green bars), which was interesting considering that depletion of these proteins strongly reduces tRNA and tricRNA production in a cellular model (12). This observation could be due to lack of transgene expression, as has been observed for other shRNA transgenes (36). Strikingly, we only observed significant adult viability defects upon depletion of RtcB ligase or other members of the ligation complex (Fig. 2, orange bars). As an additional control, we depleted *CG33057*, the putative *Drosophila* ortholog of the yeast 2′-phosphotransferase enzyme, Tpt1. Previously, we found that depletion of *CG33057/Tpt1* in a cell culture model had no effect on tricRNA formation (12). Here, we found that depletion of Tpt1 *in vivo* had no significant effect on pupal (not shown) or adult (Fig. 2, purple bars) viability. Overall, we observed a range of viability phenotypes when knocking down known tRNA processing factors in *Drosophila*.

**Fig 2.**
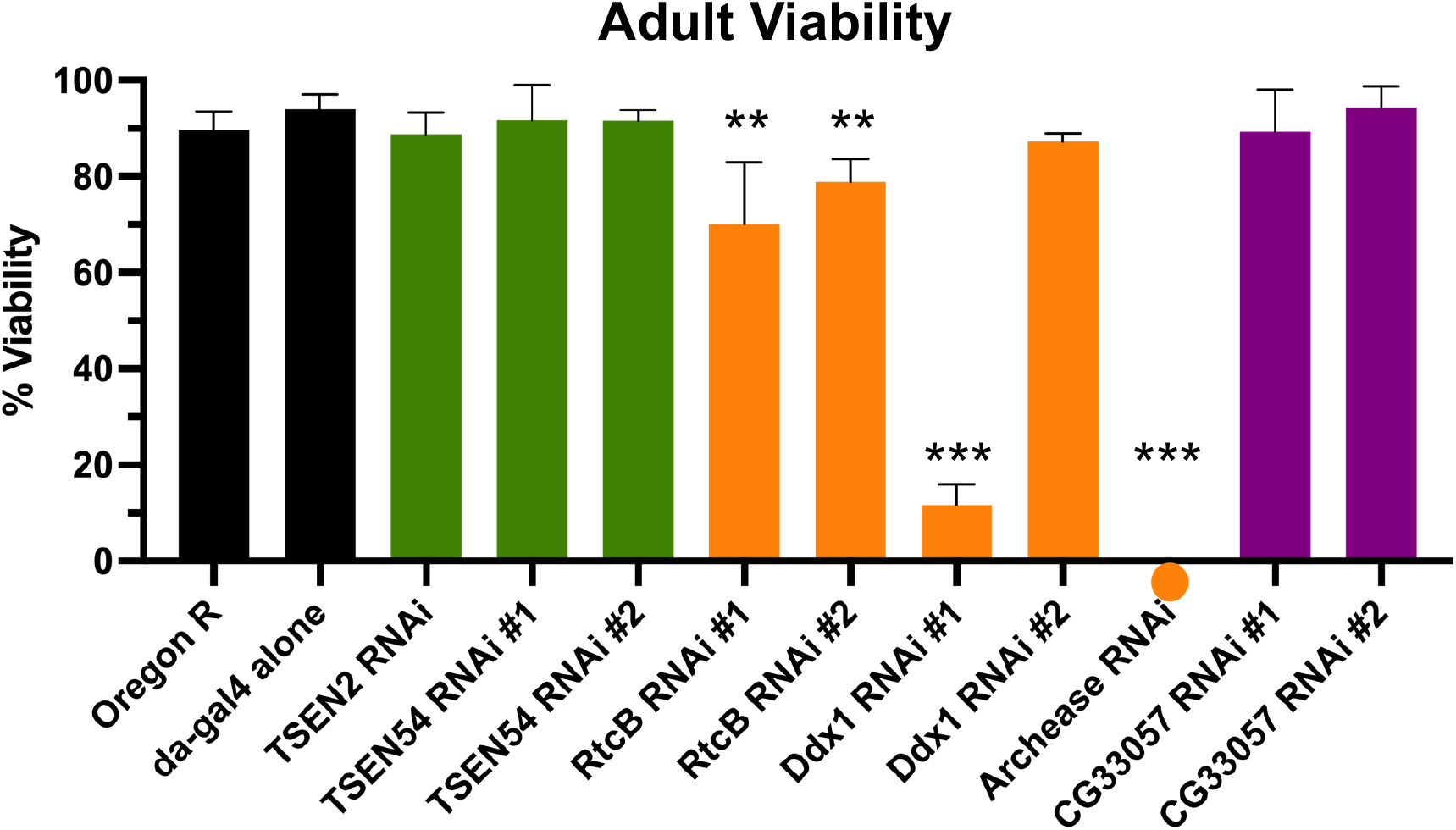
Ubiquitous depletion of *Drosophila* tRNA ligation factors, but not cleavage factors, affects viability. Adult viability of control and RNAi crosses. Oregon R and da-Gal4 alone (black) are controls. For each factor, only the da-Gal4 x UAS-RNAi cross is shown. The green bars are knockdown of pre-tRNA cleavage factors; the orange bars are knockdown of tRNA ligation factors; and the purple bars are knockdown of healing/sealing pathway factors. **p<0.01, ***p<0.001, student’s t-test. Note that a circle is placed below the x-axis indicates a zero value rather than absence of data.

### tRNA processing factor mutants exhibit larval locomotion defects

We next focused on mobility of the mutant animals. Previous reports from a Clp1-kinase dead knock-in mouse model found progressive locomotor defects in the homozygous mutants, including altered gait, reduced stride length, and impaired balance (30). Furthermore, PCH patients are often reported to have a lack of motor development (18). To determine if the *Drosophila* mutants also displayed locomotor defects, we performed assays on wandering third instar larvae. All of the homozygous mutants except for *Tsen54*^2^ displayed a significant reduction in crawling speed, measured in body lengths per second to normalize for larval size (Fig. 3B). Interestingly, the locomotion defect was not quite as severe in the cbc mutants; these animals displayed crawling speed more similar to wild-type, although still significantly reduced. Representative crawling paths can be seen in Fig. 3A. The wild-type animals crawled in a generally straight motion path, whereas the mutants’ motion paths were typically shorter and had more turns. Overall, we observed locomotion defects in most of the mutants, consistent with both animal model and patient data.

**Fig 3.**
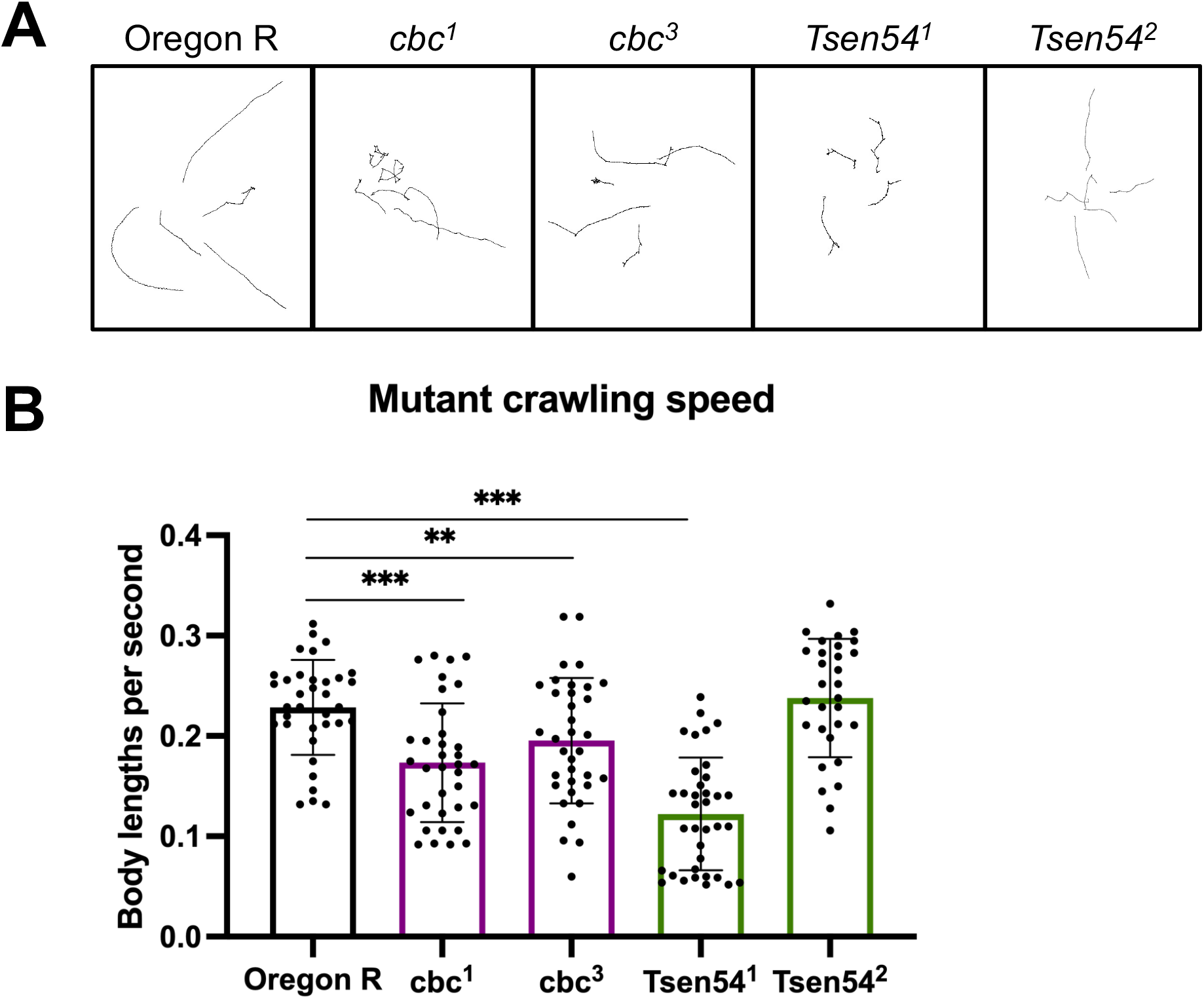
tRNA processing factor mutants exhibit locomotion defects. (A) Representative crawling paths of homozygous mutant and wild-type third instar larvae. (B) Crawling speed of homozygous mutant and wild-type third instar larvae measured in body lengths per second. **p<0.01; ***p<0.001, student’s t-test.

### tRNA processing factor mutants have reduced brain lobe size

The most prominent feature of PCH patients is microcephaly; the frequency of other phenotypes appears to be dependent on the specific disease subtype (18,21). Depletion of *Tsen54* in zebrafish embryos leads to brain hypoplasia as well as structural defects (23). Clp1 kinase-dead mice exhibited both reduced brain weight and volume, as well as structural abnormalities (30). Furthermore, zebrafish bearing a homozygous nonsense mutation in *Clp1* displayed gross head morphological defects and a massive increase in TUNEL-positive cells (24). To determine if the tRNA processing factor mutants recapitulate the brain phenotypes observed in mice, zebrafish, and human patients, we dissected third instar larval brains and performed immunofluorescence. Brains were stained with anti-HRP to detect neurons, phalloidin-FITC to detect actin, and DAPI to detect DNA. Imaging these stained brains revealed that all mutants displayed a reduction in brain lobe size (Fig. 4A, quantified in Fig. 4B), including a positive control known to exhibit microcephaly (37). We quantified brain lobe volume as described by (37) and found that the mutants and positive control showed a significant reduction in brain lobe volume.

**Fig 4.**
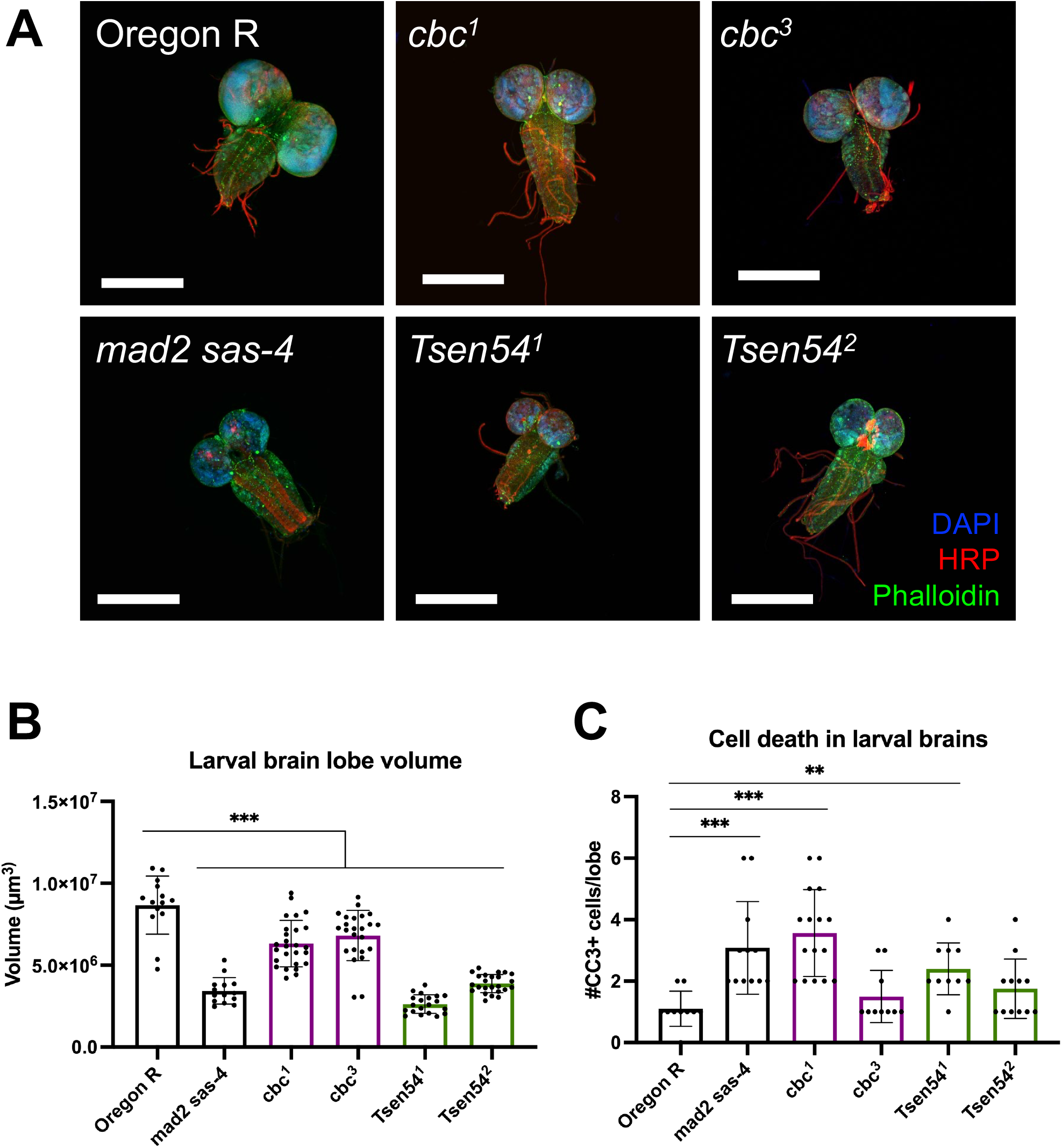
tRNA processing factor mutants exhibit locomotion defects.tRNA processing factor mutants display significantly reduced brain lobe volume. (A) Representative images of wandering third instar larval brains stained with anti-HRP, phalloidin-FITC, and DAPI. Scale bar is 300 μm. (B) Quantification of the brain lobe volume from the images in (A). The controls are in black, *cbc* mutants are in purple, and *Tsen54* mutants are in green. ***p<0.001, student’s t-test. (C) Quantification of apoptotic cells per brain lobe. **p<0.01, student’s t-test.

We next tested whether the reduced volume was due to programmed cell death. Accordingly, we stained brains with an antibody against cleaved caspase 3 (CC3) to detect apoptotic cells (Fig. 4C). The positive control *mad2 sas-*4 double mutant displayed a significant increase in the number of CC3+ brain cells per lobe. Interestingly, the stronger of the two alleles for each processing factor (*cbc*^1^ and *Tsen54*^1^) both had a significant increase in apoptotic cells; however, the weaker of the two alleles (*cbc*^3^ and *Tsen54*^2^) did not show a significant difference in apoptotic cells, despite having significantly smaller brain lobes. Taken together, these data show that all tRNA processing factor mutants have significantly smaller brains than wild-type larvae, and this reduction in volume is due, at least in part, to apoptosis in the stronger mutants.

### Tissue-specific depletion of cbc causes viability and locomotion defects

Although tRNA splicing is a ubiquitous cellular process, certain tissues such as motor neurons seem to be more sensitive to mutations in tRNA processing enzymes. As evidence of a tissue-specific need for tRNA biogenesis, the neurodegenerative phenotypes observed in the Clp1 kinase-dead mice could be rescued by expression of wild-type Clp1 only in motor neurons via the Hb9 promoter (30). To test if there is similar tissue specificity in flies, we knocked down *cbc* using various Gal4 drivers that target neurons and muscles (Table 3). As a control, we knocked down *cbc* in eye tissue (GMR-Gal4) and observed no strong defects in viability or locomotion (Fig. 5). We found the strongest developmental (Figs. 5A and B) and locomotor (Fig. 5C) defects using a combined neuron + muscle driver, C15-Gal4 (32). To further refine our analysis, we next used more specific neuron and muscle Gal4 drivers (Table 3). Depletion of *cbc* in all neurons (C155-Gal4) or motor neurons (C164-Gal4) had a strong effect on pupal and adult viability, as well as larval locomotion (Fig. 5). We observed weaker effects on viability and locomotion with glutamatergic (OK371-Gal4) or cholinergic (Cha-Gal4) neuron drivers (Fig. 5). Interestingly, although we did not observe any notable pupal viability defects using muscle drivers (C57-Gal4, Mef2-Gal4), depleting *cbc* from these tissues strongly reduced adult viability (Figs. 5A and B). Furthermore, *cbc* knockdown via the Mef2-Gal4 driver severely affected larval crawling speed (Fig. 5C). These results suggest that, consistent with mouse data, cbc protein functions in motor neurons. However, we also show here that cbc is additionally important in muscles. Further experiments can shed light on the role of cbc in both tissues.

**Fig 5.**
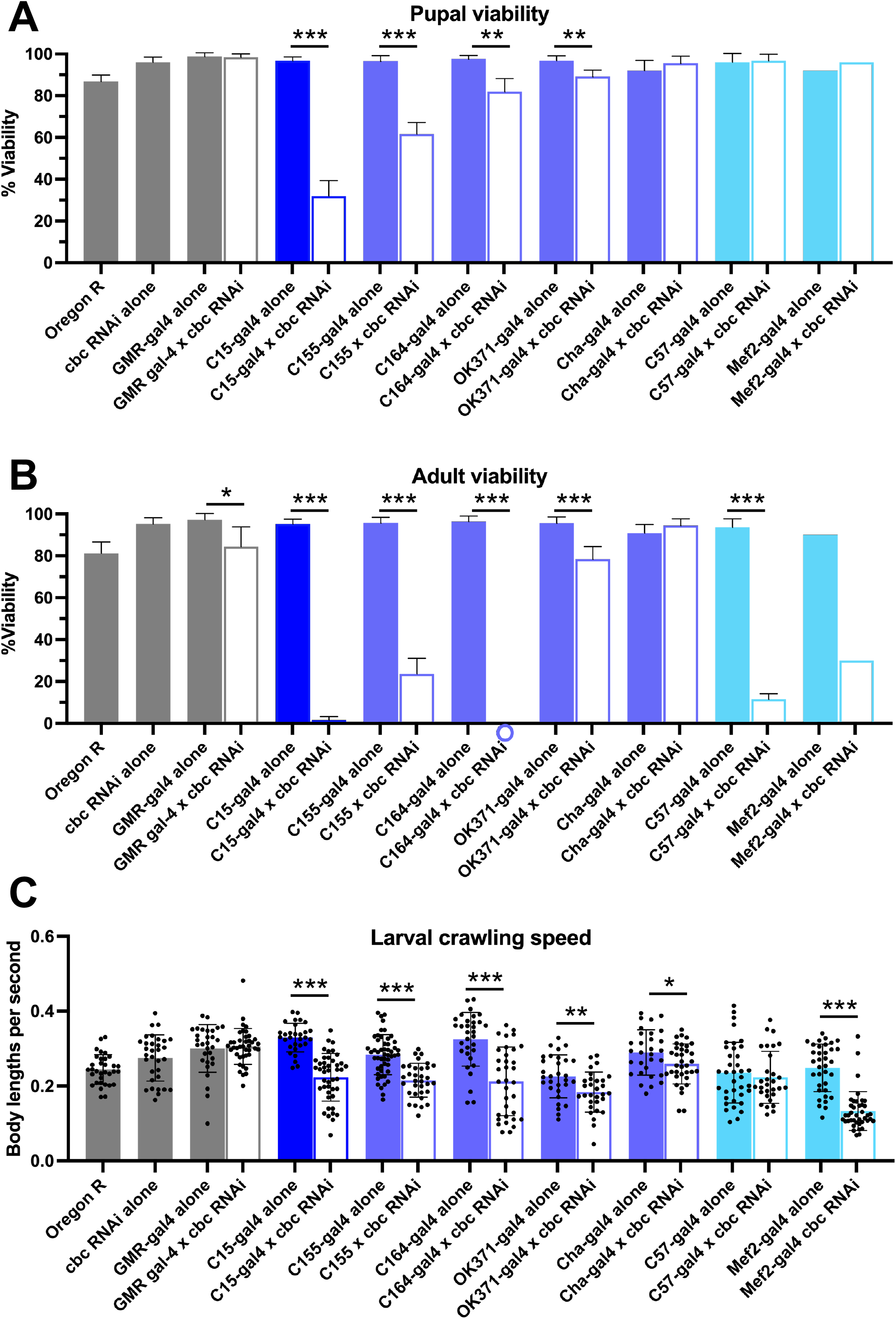
tRNA processing factor mutants exhibit locomotion defects.Evidence for a tissue-specific requirement for cbc. (A) Pupal and (B) adult viability of control and RNAi crosses. Oregon R and cbc RNAi alone are negative controls. For each cross, the shaded bar is the Gal-4 element alone, and the unshaded bar is the RNAi cross. Gray bars are controls, dark blue bars are combined neuron + muscle drivers, purple bars are neuron drivers, and light blue bars are muscle drivers. *p<0.05; **p<0.01; ***p<0.001, student’s t-test. Note that in some crosses a circle is placed below the x-axis to indicate a zero value rather than absence of data. Also note that the Mef2 results are from only one trial. (C) Crawling speed of wandering third instar larvae from control and RNAi crosses measured in body lengths per second.

## Discussion

Genetic diseases that result in the loss of neurons within the pons and cerebellum of the human brain typically do so early in development and thus were originally described as a form of underdevelopment or hypoplasia (reviewed in (17)). Subsequent studies have revealed that pontocerebellar hypoplasias are more accurately described as a heterogeneous group of neurodegenerative disorders that typically present prenatally. Curiously, at least eleven different PCH subtypes have been described, and most of them are caused by mutations in genes that encode RNA processing factors or regulators thereof (38). The sporadic occurrence and heterogeneous presentation of rare genetic diseases like PCH make them quite difficult to study. In such cases, researchers turn to model systems to better understand these disorders.

Mutations in components of the *Drosophila* RNA exosome (39) have been shown to cause PCH-like phenotypes. This work represents the first report of tRNA splicing-based models of PCH in the fruitfly. Using available *cbc* and *Tsen54* mutants, we observed disease-related phenotypes in a genetically-tractable model organism. We also utilized ubiquitous and tissue-specific RNAi to ascertain the *in vivo* function of tRNA splicing factors. Future refinements to these models should elucidate the molecular mechanisms by which the tRNA splicing machinery interfaces with metazoan neurological pathways and machineries.

It is important to note that our experiments were performed on available mutant lines from stock centers, rather than with specific patient-derived mutations. However, despite the lack of such specific mutations in our models, we observed PCH-related phenotypes in nearly all of the mutants tested. Strikingly, these animals all exhibited microcephaly (Fig. 4), similar to established PCH phenotypes (16,17,20,21). Although tRNA splicing is implicated in PCH (20,24,25), it is possible that these diseases are caused by a non-tRNA splicing function of the TSEN complex or its presumptive regulatory factor, CLP1. In this case, we would need to generate patient-derived mutations in TSEN complex members or cbc. Due to the ease of genetic engineering in *Drosophila*, such disease-specific mutations, such as the R140H mutation in *CLP1*, would in theory be straightforward to make. The assays performed above could be repeated in these mutant animals to generate a more relevant disease model.

One benefit of using *Drosophila* as a model system is that we can directly examine tricRNAs in mutant and RNAi animals. Because tRNA and tricRNA biogenesis are affected in the same way by processing factor mutations (12), tricRNA levels can be used as a proxy for tRNAs, which can sometimes be difficult to detect due to their abundance, stability, and modifications. Based on previous work (12,29), we predict that tricRNA abundance will be much lower in the *Tsen54* mutants. On the other hand, we hypothesize that the *cbc* mutants will have higher levels of tricRNAs than wild-type. If cbc indeed functions as a kinase on tRNA 3′ exons and introns *in vivo*, RtcB would be unable use these 5′-phosphorylated molecules as substrates for ligation. Thus, mutation or depletion of cbc could remove this negative regulation of tRNA and tricRNA biogenesis. Northern blots of endogenous tricRNAs will reveal if these hypotheses are true. These experiments would fit into a larger body of work on the link between tRNA processing and neurological disease, and perhaps add insight to the mechanism of disease onset and progression.

## Acknowledgements

Work in the Matera lab was supported by NIH grant R35-GM136435 (to AGM). We thank Dr. John Poulton (UNC Nephrology) for gift of the mad2, sas-4 double mutant flies, as well as for sharing the immunofluorescent brain imaging protocol. We also thank Tony Perdue (UNC Biology) for assistance with microscopy. We gratefully acknowledge Dr. Amanda Raimer and Dr. Ashlyn Spring for help with troubleshooting the locomotion assay analysis.

## References

1. Phizicky EM, Hopper AK. tRNA biology charges to the front. Genes Dev [Internet]. 2010 Sep 1 [cited 2014 Dec 2];24(17):1832–60. Available from: http://www.pubmedcentral.nih.gov/articlerender.fcgi?artid=2932967&tool=pmcentrez&rendertype=abstract

2. Chan PP, Lowe TM. GtRNAdb: A database of transfer RNA genes detected in genomic sequence. Nucleic Acids Res. 2009;37(SUPPL. 1):93–7.

3. Chan PP, Lowe TM. GtRNAdb 2.0: An expanded database of transfer RNA genes identified in complete and draft genomes. Nucleic Acids Res. 2016;44(D1):D184–9.

4. Yoshihisa T. Handling tRNA introns, archaeal way and eukaryotic way. Front Genet [Internet]. 2014 Jan [cited 2014 Oct 14];5(July):1–16. Available from: http://www.pubmedcentral.nih.gov/articlerender.fcgi?artid=4090602&tool=pmcentrez&rendertype=abstract

5. Schmidt CA, Matera AG. tRNA introns: Presence, processing, and purpose. Wiley Interdiscip Rev RNA. 2020;11(3):e1538.

6. Paushkin S, Patel M, Furia B, Peltz S, Trotta C. Identification of a human endonuclease complex reveals a link between tRNA splicing and pre-mRNA 3′ end formation. Cell [Internet]. 2004 [cited 2014 Dec 11];117:311–21. Available from: http://www.sciencedirect.com/science/article/pii/S0092867404003423

7. Trotta CR, Miao F, Arn EA, Stevens SW, Ho CK, Rauhut R, et al. The Yeast tRNA Splicing Endonuclease: A Tetrameric Enzyme with Two Active Site Subunits Homologous to the Archaeal tRNA Endonucleases. Cell [Internet]. 1997 Jun;89(6):849–58. Available from: http://linkinghub.elsevier.com/retrieve/pii/S0092867400802706

8. Abelson J, Trotta CR, Li H. tRNA Splicing. J Biol Chem [Internet]. 1998 May 22 [cited 2014 Nov 13];273(21):12685–8. Available from: http://www.jbc.org/cgi/doi/10.1074/jbc.273.21.12685

9. Popow J, Englert M, Weitzer S, Schleiffer A, Mierzwa B, Mechtler K, et al. HSPC117 is the essential subunit of a human tRNA splicing ligase complex. Science [Internet]. 2011 Feb 11 [cited 2014 Nov 13];331(6018):760–4. Available from: http://www.ncbi.nlm.nih.gov/pubmed/21311021

10. Popow J, Jurkin J, Schleiffer A, Martinez J. Analysis of orthologous groups reveals archease and DDX1 as tRNA splicing factors. Nature [Internet]. 2014 Jul 3 [cited 2014 Nov 13];511(7507):104–7. Available from: http://www.ncbi.nlm.nih.gov/pubmed/24870230

11. Lu Z, Filonov GS, Noto JJ, Schmidt CA, Hatkevich TL, Wen Y, et al. Metazoan tRNA introns generate stable circular RNAs in vivo. RNA. 2015;21(9):1554–65.

12. Schmidt CA, Giusto JD, Bao A, Hopper AK, Matera AG. Molecular determinants of metazoan tricRNA biogenesis. Nucleic Acids Res. 2019;47(12):6452–65.

13. Salgia S, Singh S, Gurha P, Gupta R. Two reactions of Haloferax volcanii RNA splicing enzymes: Joining of exons and circularization of introns. RNA [Internet]. 2003 [cited 2014 Dec 11];9:319–30. Available from: http://rnajournal.cshlp.org/content/9/3/319.short

14. Lopes RRS, Kessler AC, Polycarpo C, Alfonzo JD. Cutting, dicing, healing and sealing: the molecular surgery of tRNA. Wiley Interdiscip Rev RNA. 2015;6(3):337–49.

15. Wu J, Hopper AK. Healing for destruction: TRNA intron degradation in yeast is a two-step cytoplasmic process catalyzed by tRNA ligase Rlg1 and 5′-to-3′ exonuclease Xrn1. Genes Dev. 2014;28(14):1556–61.

16. Battini R, D’Arrigo S, Cassandrini D, Guzzetta A, Fiorillo C, Pantaleoni C, et al. Novel mutations in TSEN54 in pontocerebellar hypoplasia type 2. J Child Neurol [Internet]. 2014 Apr [cited 2014 Dec 5];29(4):520–5. Available from: http://www.ncbi.nlm.nih.gov/pubmed/23307886

17. Namavar Y, Barth PG, Kasher PR, van Ruissen F, Brockmann K, Bernert G, et al. Clinical, neuroradiological and genetic findings in pontocerebellar hypoplasia. Brain [Internet]. 2011 Jan [cited 2014 Dec 5];134(Pt 1):143–56. Available from: http://www.ncbi.nlm.nih.gov/pubmed/20952379

18. Namavar Y, Chitayat D, Barth PG, van Ruissen F, de Wissel MB, Poll-The BT, et al. TSEN54 mutations cause pontocerebellar hypoplasia type 5. Eur J Hum Genet [Internet]. 2011 Jun [cited 2014 Dec 5];19(6):724–6. Available from: http://www.pubmedcentral.nih.gov/articlerender.fcgi?artid=3110057&tool=pmcentrez&rendertype=abstract

19. Cassandrini D, Biancheri R, Tessa A, Di Rocco M, Di Capua M, Bruno C, et al. Pontocerebellar hypoplasia. Neurology. 2010;75:1459–64.

20. Breuss MW, Sultan T, James KN, Rosti RO, Scott E, Musaev D, et al. Autosomal-Recessive Mutations in the tRNA Splicing Endonuclease Subunit TSEN15 Cause Pontocerebellar Hypoplasia and Progressive Microcephaly. Am J Hum Genet [Internet]. 2016;99(1):228–35. Available from: http://linkinghub.elsevier.com/retrieve/pii/S0002929716301586

21. Budde BS, Namavar Y, Barth PG, Poll-The BT, Nürnberg G, Becker C, et al. tRNA splicing endonuclease mutations cause pontocerebellar hypoplasia. Nat Genet. 2008;40(9):1113–8.

22. Valayannopoulos V, Michot C, Rodriguez D, Hubert L, Saillour Y, Labrune P, et al. Mutations of TSEN and CASK genes are prevalent in pontocerebellar hypoplasias type 2 and 4. Brain [Internet]. 2012 Jan [cited 2014 Dec 5];135(Pt 1):1–5. Available from: http://www.ncbi.nlm.nih.gov/pubmed/21609947

23. Kasher PR, Namavar Y, van Tijn P, Fluiter K, Sizarov A, Kamermans M, et al. Impairment of the tRNA-splicing endonuclease subunit 54 (tsen54) gene causes neurological abnormalities and larval death in zebrafish models of pontocerebellar hypoplasia. Hum Mol Genet [Internet]. 2011 Apr 15 [cited 2014 Dec 5];20(8):1574–84. Available from: http://www.ncbi.nlm.nih.gov/pubmed/21273289

24. Schaffer AE, Eggens VRC, Caglayan AO, Reuter MS, Scott E, Coufal NG, et al. CLP1 founder mutation links tRNA splicing and maturation to cerebellar development and neurodegeneration. Cell [Internet]. 2014 Apr 24 [cited 2014 Nov 30];157(3):651–63. Available from: http://www.ncbi.nlm.nih.gov/pubmed/24766810

25. Karaca E, Weitzer S, Pehlivan D, Shiraishi H, Gogakos T, Hanada T, et al. Human CLP1 mutations alter tRNA biogenesis, affecting both peripheral and central nervous system function. Cell [Internet]. 2014 Apr 24 [cited 2014 Jul 9];157(3):636–50. Available from: http://www.ncbi.nlm.nih.gov/pubmed/24766809

26. Wafik M, Taylor J, Lester T, Gibbons RJ, Shears DJ. 2 new cases of pontocerebellar hypoplasia type 10 identified by whole exome sequencing in a Turkish family. Eur J Med Genet. 2018;61(5):273–9.

27. Weitzer S, Martinez J. The human RNA kinase hClp1 is active on 3’ transfer RNA exons and short interfering RNAs. Nature [Internet]. 2007 May 10 [cited 2014 Nov 13];447(7141):222–6. Available from: http://www.ncbi.nlm.nih.gov/pubmed/17495927

28. de Vries H, Ruegsegger U, Hubner W, Friedlein A, Langen H, Keller W. Human pre-mRNA cleavage factor IIm contains homologs of yeast proteins and bridges two other cleavage factors. EMBO J. 2000;19(21):5895–904.

29. Hayne CK, Schmidt CA, Haque MI, Gregory Matera A, Stanley RE. Reconstitution of the human tRNA splicing endonuclease complex: Insight into the regulation of pre-tRNA cleavage. Nucleic Acids Res. 2020;48(14):7609–22.

30. Hanada T, Weitzer S, Mair B, Bernreuther C, Wainger BJ, Ichida J, et al. CLP1 links tRNA metabolism to progressive motor-neuron loss. Nature [Internet]. 2013 Mar 28 [cited 2014 Nov 21];495(7442):474–80. Available from: http://www.pubmedcentral.nih.gov/articlerender.fcgi?artid=3674495&tool=pmcentrez&rendertype=abstract

31. Schmidt CA, Noto JJ, Filonov GS, Matera AG. A Method for Expressing and Imaging Abundant, Stable, Circular RNAs in Vivo Using tRNA Splicing. Methods Enzymol. 2016;572:215–36.

32. Spring AM, Raimer AC, Hamilton CD, Schillinger MJ, Matera AG. Comprehensive Modeling of Spinal Muscular Atrophy in Drosophila melanogaster. Front Mol Neurosci. 2019;12(May):1–16.

33. Larkin A, Marygold SJ, Antonazzo G, Attrill H, dos Santos G, Garapati P V., et al. FlyBase: Updates to the Drosophila melanogaster knowledge base. Nucleic Acids Res. 2021;49(D1):D899–907.

34. Bischof J, Björklund M, Furger E, Schertel C, Taipale J, Basler K. A versatile platform for creating a comprehensive UAS-ORFeome library in Drosophila. Dev. 2012;140(11):2434–42.

35. Brand AH, Perrimon N. Targeted gene expression as a means of altering cell fates and generating dominant phenotypes. Development [Internet]. 1993;118(2):401–15. Available from: http://www.ncbi.nlm.nih.gov/pubmed/8223268

36. Dietzl G, Chen D, Schnorrer F, Su KC, Barinova Y, Fellner M, et al. A genome-wide transgenic RNAi library for conditional gene inactivation in Drosophila. Nature. 2007;448(7150):151–6.

37. Poulton JS, Cuningham JC, Peifer M. Centrosome and spindle assembly checkpoint loss leads to neural apoptosis and reduced brain size. J Cell Biol. 2017;216(5):1255–65.

38. Van Dijk T, Baas F, Barth PG, Poll-The BT. What’s new in pontocerebellar hypoplasia? An update on genes and subtypes. Orphanet J Rare Dis. 2018;13(1):1–16.

39. Morton DJ, Jalloh B, Kim L, Kremsky I, Nair RJ, Nguyen KB, et al. A drosophila model of pontocerebellar hypoplasia reveals a critical role for the RNA exosome in neurons. PLoS Genet [Internet]. 2020;16(7):1–28. Available from: http://dx.doi.org/10.1371/journal.pgen.1008901

